# The hotspots in primate cortical brain evolution support supramodal cognitive flexibility

**DOI:** 10.1101/333930

**Authors:** Markus H. Sneve, Håkon Grydeland, Marcello G. P. Rosa, Tomáš Paus, Tristan Chaplin, Kristine Walhovd, Anders M. Fjell

## Abstract

Primate cortical evolution has been characterized by massive and disproportionate expansion of a set of specific regions in the neocortex. The associated increase in neocortical neurons comes with a high metabolic cost, thus the functions served by these regions must have conferred significant evolutionary advantage. Here, across a series of experiments, we show that the evolutionary high-expanding ‘hotspots’ – as estimated from patterns of evolutionary expansion from several primate species – share functional connections with different brain networks in a context-dependent manner. This capacity of the hotspots to connect flexibly with various specialized brain networks depending on particular cognitive requirements suggests that their selective growth and sustainment in evolution has been linked to their involvement in supramodal cognition. In accordance with an evolutionary-developmental view, we find that this ability to flexibly modulate functional connections as a function of cognitive state emerges gradually through childhood, with a prolonged developmental trajectory plateauing in young adulthood.

---

The most striking feature of the human brain when compared with brains of other primates, is its massively expanded cerebral cortex, mainly due to a higher number of cortical neurons (Herculano-Houzel 2012; Buckner and Krienen 2013). The growth of the primate cortex has not been uniform, however, with a set of ‘hotspot’ regions in lateral temporal, parietal and prefrontal cortex showing disproportionally high expansion (Hill et al. 2010; Chaplin et al. 2013). Well-supported models suggest that this non-uniform growth has followed allometric scaling laws, and that the hotspots’ massive expansion in evolution therefore is a predicted consequence of growing a bigger brain (Finlay and Darlington 1995; Toro et al. 2008; Herculano-Houzel 2012; Chaplin et al. 2013; Rilling 2014; Amlien et al. 2016; although see e.g. Smaers et al. 2017). Nevertheless, the metabolic costs associated with sustaining a higher number of neurons are high (Ringo 1991; Herculano-Houzel 2011) and impose limitations on the growth of other, non-neuronal, physical capacities such as body size (Fonseca-Azevedo and Herculano-Houzel 2012). An intriguing question is therefore: what functions do the high-expanding regions in primate brain evolution serve, which may have offset these potential natural selection costs?

Expansion hotspots refer to the cortical regions that differ the most between extant primates of different brain sizes (Chaplin et al. 2013), and are considered a useful proxy for the parts of cortex that have undergone the most expansion during evolution (Hill et al. 2010; Buckner and Krienen 2013). As prototypical members of association cortex, the evolutionary hotspots are involved in various tasks and functional systems, and have been theorized to serve relational reasoning and integrative higher-order cognition (Krienen et al. 2014; Vendetti and Bunge 2014). In support of these views, measures of general cognitive abilities in human adults have been shown to correlate positively with cortical surface-area within high-expanding regions (Fjell et al. 2015), associations not found within low-expanding cortex (Vuoksimaa et al. 2016). Furthermore, when compared with brain regions showing less evolutionary expansion, the hotspots show higher growth of the surface area during human development (Hill et al. 2010; Amlien et al. 2016), and this pattern is more pronounced in individuals with high intelligence (Schnack et al. 2015).

In the current study, we tested the hypothesis that the emergence and sustainment of the expansion hotspots have facilitated supramodal cognition (Goldman-Rakic 1988), conceptualized as the integration of information from across the brain in a flexible manner. By taking advantage of recent developments in brain network analysis (Rubinov and Sporns 2010), and applying these to histological data and magnetic resonance imaging (MRI) scans of multiple primate brains, as well as a large sample of humans during different cognitive states and at different stages in development, we were able to directly test the predictions that high-expanding cortex 1) has broad functional connections with different specialized brain networks; 2) is flexibly engaged by a diverse set of cognitive tasks; 3) is more central to communication flow in the brain during states requiring multimodal integration than during low-demand states; 4) increases functional coupling preferentially with regions engaged by the current cognitive demands. Moreover, given the correlation between morphological changes in primate brain evolution and human development, we predict that the connectivity that enables supramodal cognition develops gradually, and fully emerges relatively late in postnatal life.

## Materials and Methods

### Human subjects

RsfMRI-data were collected in 221 healthy young adults (age range 18-38, mean age 23.8; 142 females). Task-fMRI data were collected in 105 of the same participants (age range 18-38, mean age 25.3; 69 females), as well as in 46 children and adolescents (age range 6-17, mean age 12.9; 21 females). The study was approved by the Regional Ethical Committee of South Norway, and participants provided written informed consent. Participants were required to be right-handed, speak Norwegian fluently and have (corrected to) normal hearing and vision. Clinical sequences (T2-FLAIR) were inspected by a neuroradiologist and deemed free of significant injuries or conditions.

### Non-human primates

Evolutionary cortical expansion was calculated from the brains of four simian primates: marmoset (*Callithrix jacchus*), capuchin (*Cebus apella*), macaque (*Macaca mulatta*) and human. Details about the calculations have been described in a previous publication (Chaplin et al. 2013). Briefly, surface models of the cerebral cortex of the four species were registered by deforming the models to align a set of landmarks, consisting of well-established homologous cortical regions, using the CARET software package (Van Essen et al. 2001). Expansion was calculated as the change in size of each mesh polygon, and was averaged across the marmoset to capuchin, marmoset to macaque, and macaque to human deformations. All tests involving comparisons with evolutionary expansion were restricted to measures extracted from right-hemispheric nodes.

### Experimental design

RsfMRI was collected during eyes-closed rest. The participants were instructed to not fall asleep, and confirmed compliance following the scan. During the fMRI-task, participants sequentially viewed 100 line drawings of objects and immediately produced a motor response indicating whether the object was congruent with a spoken action (either “Can you eat it” or “Can you lift it”; SFig. 2). The task is described in detail elsewhere (Sneve et al. 2015).

### MRI acquisition

Imaging was performed at a Siemens Skyra 3T MRI unit with a 24-channel head coil. For the fMRI scans (rest and task), 43 slices (transversal, no gap) were measured using T2* BOLD EPI (TR=2390 ms; TE=30 ms; flip angle=90°; voxel size=3×3×3 mm; FOV=224×224; interleaved acquisition; GRAPPA=2). The rsfMRI run produced 150 volumes and lasted ≈6 min. The task data were collected over 2 runs, each consisting of 131 volumes and lasting ≈5.2 min. Three dummy volumes were collected at the start of each fMRI scan to avoid T1 saturation effects in the analyzed data. A standard double-echo gradient-echo field map was acquired for distortion correction of the EPI images. Anatomical T1-weighted MPRAGE images consisting of 176 sagittally oriented slices were obtained using a turbo field echo pulse sequence (TR=2300 ms, TE=2.98 ms, flip angle=8°, voxel size=1×1×1 mm, FOV=256×256 mm).

### MRI preprocessing

Cortical reconstruction of the T1-weighted scans was performed with Freesurfer 5.3’s recon-all routines, and included surface inflation (Fischl et al., 1999a) and registration to a spherical atlas which utilized individual cortical folding patterns to match cortical geometry across subjects (Fischl et al., 1999b). FMRI-data were corrected for B0 inhomogeneity, motion and slice timing corrected, and smoothed (5mm FWHM) in volume space using FSL (http://fsl.fmrib.ox.ac.uk/fsl/fslwiki). Next, FMRIB’s ICA-based Xnoiseifier (FIX; Salimi-Khorshidi et al. 2014) was used to auto-classify noise components and remove them from the fMRI data. Different classifiers were used for rsfMRI and task-fMRI data. Classifiers were trained on scanner-specific datasets in which rsfMRI/task-fMRI data from 16 participants had been manually classified into signal and noise components (fMRI acquisition parameters identical to the current study). Motion confounds (24 parameters) were regressed out of the fMRI data as a part of the FIX routines. Freesurfer-defined individually estimated anatomical masks of cerebral white matter (WM) and cerebrospinal fluid / lateral ventricles (CSF) were resampled to each individual’s functional space. Following FIX, average time series were extracted from functional WM- and CSF-voxels, and were regressed out of the FIX-cleaned 4D volume. Following recent recommendations (Hallquist et al. 2013) we band-pass filtered the rsfMRI data (.009-.08Hz) after regression of confound variables. Task-fMRI data were detrended and highpass filtered with a.01Hz cutoff.

### Network analysis of rsfMRI data

A custom cortical parcellation was created in Freesurfer’s average surface space (fsaverage), using a modified N-cut algorithm (Craddock et al. 2012), consisting of 340 (170 per hemisphere) spatially contiguous, approximately equally sized nodes covering the entire cerebral cortex. The parcellation was resampled into each participant’s functional volume space using a projection factor of 0.5, i.e., half way into the cortical sheet. For each participant, we extracted mean pre-processed rsfMRI time series from all nodes and calculated a 340×340 connectivity matrix consisting of the Pearson’s *r* correlations between nodal time series. Next, we Fisher-transformed all participants’ connectivity matrices, and averaged across participants to create a “grand average” connectivity matrix on which network analysis was performed. The grand average connectivity matrix was thresholded at 5, 6, 7, 8, 9, 10, 15, and 20% edge densities. For all thresholded weighted graphs, the optimal modular resolution parameter (gamma) promoting stable decomposition results, was calculated using the Versatility approach (Shinn et al. 2017; see SFig1). Next, modular decomposition was performed with the Louvain algorithm (Blondel et al. 2008) as implemented in the Brain Connectivity Toolbox (BCT) (Rubinov and Sporns 2010), and consensus clustering (Sporns and Betzel 2016). Briefly, this involved calculating an agreement matrix from 10000 independent Louvain partitions, thresholding this empirical agreement matrix by the maximum agreement observed over 10000 randomly generated null association matrices and running clustering on the thresholded empirical agreement matrix. In the case of singleton partitions, i.e., network modules consisting of one node only – typically consisting of low signal-to-noise regions such as the temporal pole and orbitofrontal cortex – these modules were excluded from the remaining network analyses. Finally, we used the thresholded graphs’ optimal community structures to calculate two nodal network measures per graph: *participation coefficient* and *within-module degree*, representing a node’s intermodular and intramodular centrality, respectively (Rubinov and Sporns 2010). Two additional measures not requiring information about the underlying community structures were also calculated: *betweenness centrality*, representing the fraction of all shortest paths in the network that contains a given node, and *strength*, the sum of a given node’s connectivity weights to every other node (Rubinov and Sporns 2010).

### Community density analyses

To investigate high-expanding nodes’ anatomical centrality, we used a recent parcellation of human cortical intrinsic connectivity into 17 canonical networks, estimated from 1000 participants (Yeo et al. 2011). First, we extracted MNI-coordinates for every vertex in the right hemisphere in a downsampled Freesurfer surface representation (fsaverage5; 10242 vertices). Next, for each vertex, we counted the number of canonical networks present within a radius of 5, 10, 15, 20, 25, and 30mm (Euclidean distance). For each radius, the number of networks at every vertex was normalized (0-1) by the maximum number of networks found across all vertices (Power et al. 2013). Finally, the normalized community density values were averaged across radii at each vertex and correlated with evolutionary expansion estimates at the same locations.

### Flexibility analyses

The functional flexibility of each vertex on the cortical surface of the right hemisphere was estimated from publicly available data (https://surfer.nmr.mgh.harvard.edu/fswiki/BrainmapOntology_Yeo2015). Here, an author-topic hierarchical Bayesian model was used to classify 10449 experimental contrasts from fMRI experiments found in the BrainMap database (Fox and Lancaster 2002) into 12 underlying cognitive components and corresponding brain activity patterns (Yeo et al. 2015). Flexibility was defined as the number of cognitive components activating a voxel. To account for non-integer values due to projections from volume to surface space we rounded surface flexibility estimates to the nearest integer.

### Task-state connectivity

Following preprocessing, volumetric task-fMRI data were brought to fsaverage surface space. Here, for each participant, we extracted mean BOLD time series from all nodes in the custom 340-node cortical parcellation. Next, psychophysiological interaction (PPI) terms representing nodal task-related activity modulations were calculated using the generalized PPI-toolbox for Matlab (McLaren et al. 2012). For each node, this involved: 1) deconvolving the mean BOLD time series into estimates of neural events (Gitelman et al. 2003); 2) setting up a task-regressor representing the combined auditory-visual-motor-event (2s trial duration, 50 trials per fMRI run) 3) convolving the product of step 1 and 2 with a canonical hemodynamic response function (cHRF). Finally, to establish task-related functional connectivity between nodes, we estimated pairwise interactions between all nodes’ PPI-terms (concatenated over runs) using partial correlations. For each pairwise correlation, we controlled for background noise and task stimulation effects using the nodes’ mean BOLD time series and the cHRF-convolved task-regressor, respectively. This “correlational PPI” approach has been described in detail elsewhere (Fornito et al. 2012).

### State-dependent coupling analyses

105 adult participants were represented with 340×340 connectivity matrices from a resting-state and a task-state. First, the individual connectivity matrices were thresholded to only contain edges surviving FDR-correction (q<.05). Next, to allow comparison of connectivity data from different states, edge-wise connectivity weights were normalized by the average weight in the matrix (Opsahl et al. 2010). After mapping from weights to lengths (inversing the connection-weights matrices), shortest path lengths were calculated using Dijkstra’s algorithm (Dijkstra 1959). A node’s closeness centrality was calculated as the inverse of the average of its shortest path length to every other node.

### Correction for Euclidean distance

Euclidean distance between two nodes was calculated as the average distance in mm between the locations of their constituent vertices converted to MNI305 space. To correct the connectivity matrices, the Euclidean distance matrix was normalized to fall between 0 and 1 and multiplied, element-by-element, with the connectivity matrices across states and participants.

### Analyses of the developmental sample

The developmental sample consisted of the 105 adult participants described in the section “State-dependent coupling analyses” and 46 participants below 18 years of age (see “Human subjects” section). All non-adult participants were preprocessed and analyzed as described for the adult sample. One participant (age 9.3 years) was excluded from the sample due to high levels of motion (mean absolute motion over two task runs > 1.5mm). A significant positive Pearson correlation (*r =.23, p =.005)* was found in the remaining developmental sample (N=150) between estimated motion (mean absolute motion over two task runs) during the task-state and global closeness centrality (average closeness centrality over all 170 nodes, corresponding to the graph theoretical measure global efficiency (Rubinov and Sporns 2010)). A similar positive relationship between levels of motion and global closeness centrality estimates was found when investigating the adult sample in isolation (*p* =.04), but not in development sample (p >.40). The positive correlation between subject motion and global closeness centrality indicated that participants with relatively high levels of motion (which is a charateristic of young samples (Satterthwaite et al. 2012)) also tended to show high closeness centrality estimates. To allow for comparisons of closeness centrality estimates across age groups, unbiased by motion, we therefore standardized (z-scored) the estimates on a within-subject basis before comparing relative coupling differences across groups. Scores on the matrix reasoning and vocabulary subtests of the Wechsler’s Abbreviated Scale of Intelligence (WASI; Wechsler 1999) were available from 140 of the participants in the developmental sample (age range 7-38 years). Principal component analysis was run on the raw scores of the two subtests, and the first component, which explained 95.5% of the total variance, was used as a representative measure of general intelligence across participants.

### Statistical analyses

Details about quantification and the statistical analyses run are presented in the figures and the associated figure legends. The specific statistical tests used were chosen after the following considerations:

For data in Fig. 1b&c, nonparametric Spearman correlations were calculated to test for monotonic relationships between variables due to non-normal distributions (assayed using Q-Q plots). A nonparametric Kruskal-Wallis was used to compare central tendencies in evolutionary expansion across flexibility group (Fig. 1d) due to unequal variances across groups (significant Bartlett’s test: χ^2^ (8) = 3549, *p* < 1e-10)

**Figure 1.**
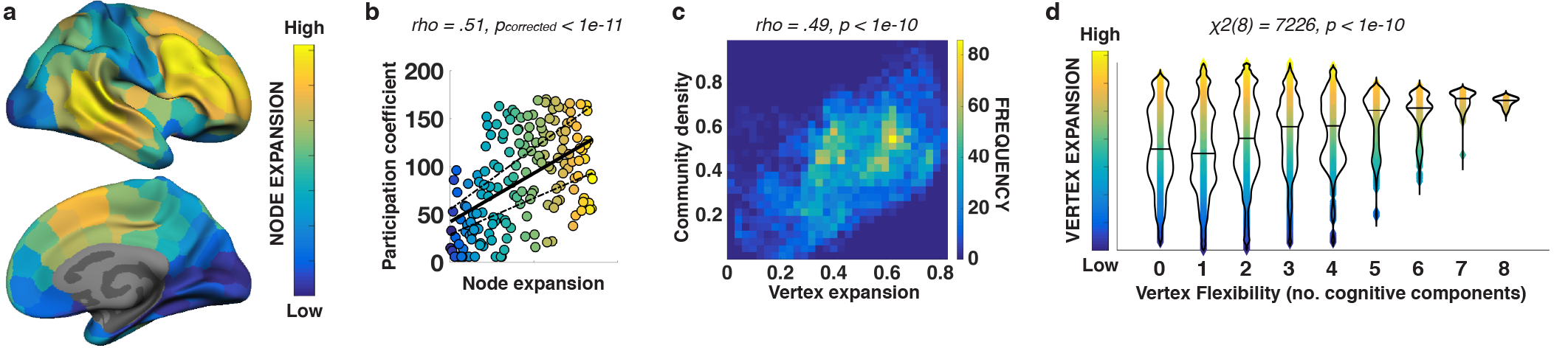
High-expanding cortex has broad functional connections and is flexibly engaged across different cognitive tasks. (a) Estimates of evolutionary expansion for 170 similarly sized nodes covering the right hemisphere. (b) Spearman correlation between node expansion and e participation coefficient. The presented values are averaged across thresholds and ranked – plotted linear relationship represent the spearman correlation. Traditional hub-measures not incorporating integrative aspects of nodal communication (within-module degree, node strength), did not show any relationships with node expansion (SFig. 1c). (c) Density plot showing the Spearman correlation between vertices’ normalized community density (calculated over 5-30mm radius in 5mm steps and averaged) and estimated evolutionary expansion(d)Violin plots showing the relationship between cortical expansion and cognitive flexibility at the vertex level. Median expansion is shown as horizontal lines. A Kruskal-Wallis test confirmed that expansion differed across flexibility groups. Post-hoc pairwise comparisons demonstrated that cortical locations involved in five or more cognitive components were more expanded than locations involved in four or components (p<1.25e-07). All reported p-values are corrected for multiple tests.

In Fig. 2a, a repeated measures ANOVA was run with two within-subject factors: “expansion bin” (5 levels) and “state” (2 levels). The reported “expansion bin x state” interaction was Greenhouse-Geisser corrected following a signficant Mauchly’s test of sphericity: χ^2^ (9) = 129, *p* < 1e-10). In Fig. 2a-d, all reported *p*-values following multiple comparisons have been corrected using Benjamini & Yekutieli’s method for controlling the False Discovery Rate (FDR) (Yekutieli and Benjamini 2001).

**Figure 2.**
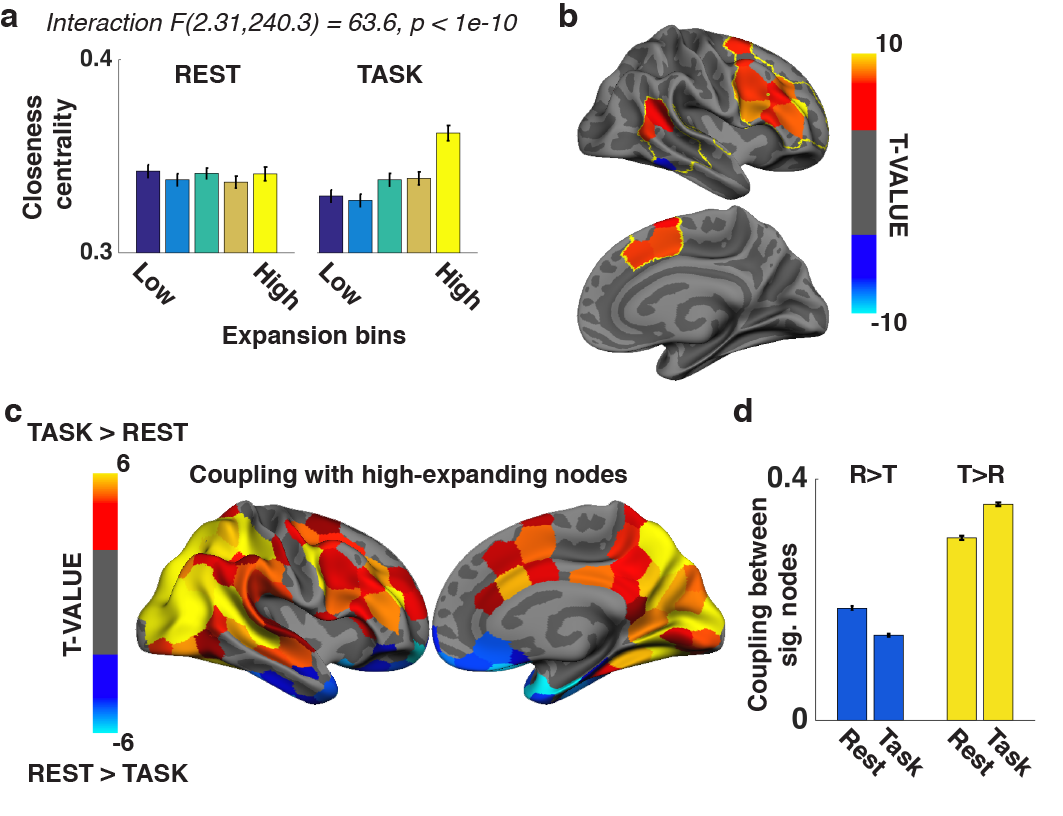
High-expanding cortex communicates differently depending on the current cognitive demands, and preferentially with regions engaged by those demands. Cortical nodes were binned per evolutionary expansion (SFig. 4). We considered the 20% highest-expanding nodes ‘hotspot’ regions, and this definition revealed three separate clusters commonly discussed in the literature (Hill et al. 2010; Chaplin et al. 2013). (a) Average closeness centrality for five expansion bins during resting- and task-state (dark blue: 20% least expanding nodes, yellow: expansion hotspots; intermediate colors: intermediate expansion levels). Following a significant “expansion bin X state” repeated measures ANOVA, paired-samples t-tests showed significant increase in closeness centrality in task-compared to resting-state for the highest-expanding nodes only: t(104)=4.27, p_FDR_ < 5.0e-04. (b) Nodes within hotspot regions (indicated by yellow lines) showing significant (paired-samples t-tests, p_FDR_ <.05) changes in closeness centrality from rest to task. (c) Nodes showing significant change (paired-samples t-tests, p_FDR_ <.05) in functional coupling (inverse of shortest path length) with expansion hotspots from rest to task across participants. (d) Mean coupling between nodes showing negative (blue) / positive effect (yellow) in panel c. Nodes falling within the hotspot regions were excluded when calculating the mean. Nodes more strongly coupled with hotspots during rest (R>T) showed higher coupling with each other during rest than during task: paired-samples t-test, t(104)=-10.56, p_FDR_ < 3.7e-18. The opposite effect was observed between nodes more strongly coupled with hotspots during the task-state (T>R): paired-samples t-test, t(104)=13.25, p_FDR_ < 4.4e-24. The presented data have been corrected for Euclidean distance between nodes. Uncorrected data show similar effects (SFig. 5). Error bars represent SEM.

In Fig. 3a, a repeated measures ANOVA was run with one within-subject factor: “expansion bin” (5 levels), and one between-subject factor: “age group” (2 levels), including subject motion (mean absolute motion over two task runs) as a covariate. The reported “expansion bin x age group” interaction was Greenhouse-Geisser corrected following a signficant Mauchly’s test of sphericity: χ^2^ (9) = 63.4, *p* < 4e-10). Nonparametric Wilcoxon rank sum tests were used for post-hoc testing due to unequal variances across groups (Bartlett’s test: χ^2^ (1) = 20.8, *p* < 6e-06). Significance of the post hoc tests was assessed following FDR-correction for multiple comparisons. In investigating the correlation between individual centrality-expansion resemblance (y-axis in Fig. 3b) and general intelligence (first principal component of raw scores from two WASI subtests), partial Spearman correlations were used with age as covariate to control for nonlinear relationships between the correlated variables and age.

**Figure 3.**
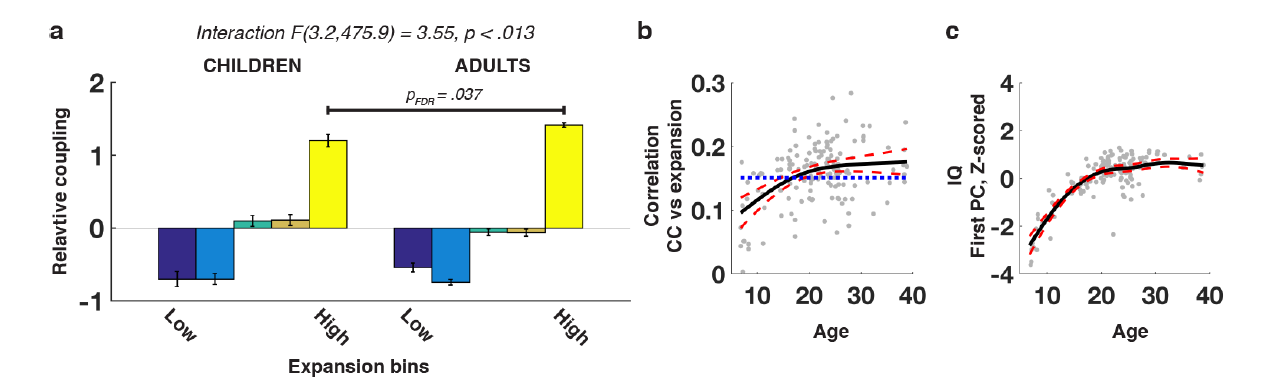
Human development of neocortical multimodal coupling patterns follows evolutionary expansion trajectories. (a) To compare hotspot’s coupling across age groups, we standardized individual closeness centrality measures across nodes into units of standard deviation. This step was performed to account for higher levels of motion in the younger participants, which correlated positively with individual estimates of global efficiency (i.e., average closeness centrality) and thus made comparisons of absolute coupling values between groups difficult. The hotspots showed stronger relative coupling than lower-expanding regions in both age groups (children: <u>t</u>(44)>8.09, < 3.0e-10; adults: t(104)>23.73, p_FDR_ < 1.0e-10). Following the significant “age group x expansion bin” interaction, post hoc Wilcoxon sum tests showed lower relative coupling of high-expanding nodes in the development sample compared to the adults (Z = −2.68, p_FDR_ < 0.37). Lower-expanding nodes did not show significant differences in relative coupling between age groups (p_FDR_ >.093). Error bars represent SEM. (b) Individual correlation between nodal closeness centrality and evolutionary expansion plotted as a function of age. The k line represents the best fitting smoothing spline (minimizing the Bayesian information criterion, BIC). Red lines represent the strapped 95% confidence interval of the fit. Blue dotted line shows the correlation coefficient at which a correlation with 169 degrees freedom is significant at p <.05 (abs(rho) >.151). Note that the BIC of a linear fit was 8.7 higher than the BIC for the optimal smoothing e, indicating that the depicted age trajectories are curvelinear. (c) First principal component calculated from raw scores on two WASI ests plotted as a function of age. Optimal fit estimated as in Fig. 3b. Spearman correlations revealed significant relationships between measure of general intelligence and individual differences in closeness centrality-vs-expansion correlation (i.e. correlating datapoints in 3b and 3c: Spearman’s rho =.32, p < 9.4e-5). Importantly, this relationship remained when controlling for nonlinear influences of age g partial Spearman correlations (rho =.18, p =.037).

Across analyses, all performed tests were two-tailed.

## Results

### High-expanding cortex has broad functional connections and is engaged flexibly across different cognitive tasks

First, we tested whether high-expanding nodes communicate more broadly across human brain networks than low-expanding nodes. Investigations in the macaque monkey have demonstrated strong positive relationships between dendritic complexity at the microscale neuronal level, and broadness of cortico-cortical neuronal connectivity profiles at the macroscale network level (Scholtens et al. 2014). Therefore, we tested whether high-expanding nodes are characterized by high *participation coefficients*, a graph theoretical measure that quantifies the degree to which a node participates in many of the brain’s subnetworks (Guimerà and Nunes Amaral 2005). Functional connectivity between nodes covering the entire cerebral cortex was calculated from resting-state functional MRI (rsfMRI) data from 221 healthy young adults. We estimated optimal community structures, i.e. the brain’s subnetworks, through modular decomposition of the group-averaged connectivity graph thresholded at different edge densities (SFig. 1a&b). Next, for each threshold, we calculated every node’s participation coefficient – a high value indicating that it communicates broadly and outside its own community. Finally, we extracted average nodal expansion between three non-human simian primates and humans from estimates of evolutionary cortical scaling (Chaplin et al. 2013) (Fig. 1a). A positive relationship was found between nodal participation coefficient values and estimates of evolutionary expansion at all edge densities (Fig. 1b and SFig. 1c), suggesting that high-expanding regions function as integrative connector nodes in the information flow between more specialized modules in human cortex (Power et al. 2013). In line with this observation, nodal *betweenness centrality* also correlated positively with expansion (SFig. 1c), demonstrating that high-expansion regions often participate in the shortest, most efficient, path between any two other nodes in the brain network (Rubinov and Sporns 2010).

Next, we tested whether the topologically broad and central connectivity profiles of high-expanding nodes – as indicated by high participation coefficient and betweenness centrality, respectively – were reflected in their anatomical centrality relative to the canonical functional networks of the human brain (Yeo et al. 2011). The measure *community density* represents the number of different networks present within a given radius from a cortical location (Power et al. 2013). We observed a positive relationship between local surface expansion and community density, indicating that high-expanding parts of the cortex have access to multiple networks present in their immediate vicinity (Fig. 1c). This closeness, both at the anatomical and the network topology-level, between high-expanding parts of the cortex and the brain’s different networks, makes high-expanding cortex ideally situated to engage in a variety of cognitive processes. To test this hypothesis, we took advantage of recent work on the BrainMap database, in which data from ≈10.000 fMRI-experiments have been merged to establish brain activity patterns common to specific types of tasks (Yeo et al. 2015). We grouped cortical surface locations based on the number of task-types (“cognitive components”) they were associated with, and then compared expansion across these levels of cognitive flexibility. In line with our hypothesis, highly flexible nodes were found predominantly in high-expanding cortex (Fig. 1d).

### High-expanding cortex communicates preferentially with regions engaged by the current cognitive demands

The above findings suggest that a key role of high-expanding cortical regions may be integration of different cognitive processes. To test this proposal directly, we estimated the *closeness centrality* of cortical nodes in 105 participants scanned using fMRI during two states: unconstrained rest, and a task-state requiring audio-visuo-motor processing (SFig. 2). Closeness centrality represents the average shortest path-length from one node to all other nodes in a network, and thus indicates how tight the functional coupling of a node is to the rest of the network (Rubinov and Sporns 2010). Expansion hotspot regions showed higher closeness centrality during the task-state than during rest and were also more tightly coupled to the rest of the network than lower-expanding nodes (Fig. 2a). Critically, this was also true when accounting for physical distance between nodes, demonstrating that the tight coupling of high-expanding nodes to the rest of the network during multimodal integration is independent of their physical locations on the cortical surface (Liu et al. 2014) (S3). The higher closeness centrality during the task-state was distributed across all expansion hotspot regions (Fig. 2b), suggesting that stronger functional coupling during effortful task-operations is a general property of high-expanding cortex and not driven by a subset of cortical regions.

To test the key proposal that hotspot regions play a central role in *supramodal* cognition – and thus interact flexibly with different parts of cortex depending on particular cognitive demands – we calculated functional coupling change (resting-state to task-state) between the expansion hotspots and all cortical nodes. During the multimodal task-state, hotspot coupling increased (i.e. path length decreased) most prominently with posterior visual perceptual regions, auditory cortex and motor cortex (Fig. 2c). In support of the hypothesis that high-expanding cortex connects with regions engaged during a given cognitive state, these regions also showed upregulated functional coupling between themselves during the task state when compared with rest (Fig. 2d). Critically, medial temporal cortex and ventromedial prefrontal cortex, regions found to be involved in memory consolidation processes during offline rest (van Kesteren et al. 2010; Euston et al. 2012), showed the opposite pattern: stronger hotspot coupling during rest than during the task state (Fig. 2c). Moreover, and in direct accordance with the proposal that the expansion hotspots interact flexibly with regions engaged in a given cognitive state, coupling between these regions was upregulated during rest when compared with the task-state (Fig. 2d).

### Human development of neocortical functional coupling patterns follows evolutionary expansion trajectories

Recent investigations of neocortical morphometry have found similarities between cortical expansion in human development and primate evolution (Fjell et al. 2015). Evolutionary high-expanding cortex, in particular, shows protracted development, and undergoes larger increases in surface area between infancy and adulthood than lower-expanding regions (Hill et al. 2010; Amlien et al. 2016). Notably, this pattern of surface change during development is more pronounced in high-intelligence samples (Schnack et al. 2015), lending support to the idea that similar developmental and evolutionary trajectories of neocortical change – albeit at very different scales – may promote the same phenotypic characteristics of higher intellectual abilities and supramodal cognition.

To test whether such correspondence exists between neocortical evolution and functional supramodal cognition characteristics in human cortical development, we collected task-state fMRI data from 46 children and adolescents (6-17 years of age, one excluded due to excessive motion), and compared closeness centrality estimates from regions differing in evolutionary expansion. As found in the adult sample, the hotspots’ closeness centrality was higher during the multimodal task state also in the developmental sample when compared with lower-expanding parts of cortex (Fig. 3a). Additionally, an interaction was observed between age group and regional expansion, indicating that the relative coupling differences between higher- and lower-expanding regions change during development. Specifically, the expansion hotspots’ closeness centrality relative to the typical (average) centrality across all nodes, i.e. their relative coupling, was found to be less developed in the young sample when compared with the adults (Fig. 3a). Importantly, less-expanding regions did not show significant differences in coupling between the two age groups. This observation fits well with recent reports of protracted surface area development of high-expanding cortex, reaching maximum expansion in adolescence (Amlien et al. 2016; Walhovd et al. 2016), and suggests that the hotspots’ roles as multimodal integrating hubs follow related developmental trajectories.

Finally, if evolutionary factors have shaped ontogenetic cortical development, we would expect the mature human brain to reflect the changes that have occurred in primate evolution to a higher degree than the immature brain (e.g. Rakic 2009). For all 150 participants in the adult and development sample, we calculated task-state closeness centrality for each of the 170 nodes in the custom neocortical parcellation and correlated these with evolutionary expansion estimates for the same nodes. Next, we fitted a nonparametric local smoothing model (Fjell et al. 2010) to delineate the age trajectory of the relationship between closeness centrality and evolutionary expansion. The relationship was found to be not significant until approximately eighteen years of age, at which point the fit revealed positive correlations between neocortical coupling pattern and expansion (Fig. 3b). Interestingly, and in line with previous morphometric reports (Fjell et al. 2015; Schnack et al. 2015), participants showing higher similarity between their nodal centrality maps and the evolutionary expansion map were characterized by higher scores on measures of general intelligence (Fig. 3c). This suggests that evolutionary concepts resonating in the functional coupling of the human neocortex are relevant for the characteristic human phenotype of higher intellectual function.

## Discussion

Our findings point to a central role of evolutionary high-expanding cortex in integrative operations during a variety of cognitive states. The functional signature of such supramodal cognition matures throughout childhood, suggesting that the hotspots develop their characteristic of broad functional connections to many of the brain’s networks in tandem with the emergence and refinement of central human cognitive skills. Their postulated overarching function in facilitating supramodal and flexible cognition is supported by the reported links between individual differences in hotspot surface area and general intelligence (Fjell et al. 2015; Schnack et al. 2015). Moreover, the functional connectivity ‘fingerprints’ of high-expanding cortex have been shown to be highly variable from participant to participant, and this characteristic overlaps well with the degree to which a brain region can be used to predict performance during different types of cognition (Mueller et al. 2013). The hotspots’ underlying cellular machinery appears optimized to support such supramodal processes: neuromorphological investigations of primate cortex have found more elaborate dendritic layouts in high-expanding compared with low-expanding regions (Elston et al. 1999), and these structural properties are more pronounced in humans than in non-human primates (Bianchi et al. 2013; Geschwind and Rakic 2013; Donahue et al. 2018).

In the current study, we interpret cortical regions demonstrating the capacity to integrate information from across the brain in a flexible manner as to take part in supramodal cognition. The claim that certain parts of the cortex, and in particular prefrontal regions, play such flexible integrative roles is hardly new (e.g.,Goldman-Rakic 1988), however we believe there is novelty in linking these broad functions to evolutionary morphological changes and their candidate behavioral phenotypes. Our results suggest that, when analyzed as a unit, high-expanding cortex shows both integrative and supramodal characteristics. However, the regions constituting the ‘hotspots’ are spread in a partly discontinuous manner across large portions of the cortex, and may show different characteristics and specializations when investigated in a more fine-grained fashion (Chaplin et al. 2013). A fascinating venue for further research could be to collect data on unimodal in addition to multimodal tasks in an attempt to disentangle modality-specific from integrative functions. Specifically, regions showing increased task-related recruitment and/or functional coupling during multimodal states when compared to unimodal states could be said to be integrative, while regions demonstrating differential coupling patterns over varying task-requirements would support supramodal flexibility. The current results suggest that one would find regions fulfilling both criteria primarily within evolutionary high-expanding cortex.

While functional connectivity measures are reflective of underlying anatomical connectivity (Vincent et al. 2007; Honey et al. 2009; Hermundstad et al. 2013), they are nevertheless estimated from covariations in signal time-series and can thus be affected by mechanisms other that direct interactions between neuronal populations. In the current study, we do not base any conclusions on observations from single edges (i.e., simple bivariate correlations between two regions), but rely on graph theoretical nodal summary measures – such as closeness centrality – and regions’ relative roles in the brain network along these measures. Moreover, due to the within-subject nature of our coupling analyses, we base our conclusions about hotspot supramodality on the observed consistent *modulations* in functional coupling across states, not the presence/absence of specific connections. Finally, while based on functional connectivity, our claims are strengthened by the observed consistencies across modalities (Community Density analyses) and with independent datasets (Flexibility analyses), as well as by the replication of selective increase in functional hotspot coupling during multimodal requirements across two independent groups of participants (Developmental Sample analyses).

The present study is based on interpretation of functional data obtained in humans, correlated with estimates of the differential expansion of parts of the cerebral cortex in primate evolution. To date, these estimates have been derived from careful histological reconstruction of single individual brains from different species, followed by computational registration of 3-dimensional models (Chaplin et al. 2013). The reliance of these estimates on individual brains, rather than population averages, represents a possible limitation of the precision of the present analyses. It should be noted, however, that the differences in expansion that underlie our conclusions are very substantial, relative to the likely degree of individual variation, or errors derived from incorrect assignment of cytoarchitectural boundaries. For example, the differential expansion between the most notable hotspots (temporoparietal junction, ventrolateral prefrontal cortex, and dorsal anterior cingulate cortex) and other regions, such as the primary visual cortex and parahippocampal gyrus, is as high as 16-fold between macaque and human (Hill et al. 2010), which is well beyond the variation observed in the volumes of cytoarchitectural areas between individuals of a single primate species (cf. Majka et al. 2016; Woodward et al. 2018). Moreover, although quantitative details of the expansion of different cortical nodes vary slightly depending on which species are used for comparison, the locations of the hotspots are remarkably stable (Chaplin et al. 2013). Finally, although some controversy remains about the relative expansion of some regions of the cerebral cortex in primate evolution (e.g. frontal lobe; Barton and Venditti 2013; Sherwood and Smaers 2013), it must be noted that the estimates used in the present study are based on quantitative analyses of species to species registration that took into consideration cytoarchitectural boundaries of areas that are well-defined, rather than gross morphological features of the brain; recent studies that also used cytoarchitecture to guide registration have confirmed that differential expansion exists (Mansouri et al. 2017; Donahue et al. 2018).

In conclusion, the present results add to existing knowledge by showing how the hotspot regions change in their role as communication hubs in different cognitive states, and that this power of the hotspot regions gradually emerges during childhood and adolescence development.

## Acknowledgements

We would like to thank Ed Bullmore (University of Cambridge) for helpful comments on an earlier draft of the manuscript, and Didac Vidal Piñeiro (University of Oslo) for help with analyses. The authors declare no competing interests.

